# Silencing and stimulating the medial amygdala impairs ejaculation but not sexual incentive motivation in male rats

**DOI:** 10.1101/2021.01.07.425687

**Authors:** Patty T. Huijgens, Roy Heijkoop, Eelke M.S. Snoeren

## Abstract

The medial amygdala (MeA) is a sexually dimorphic brain region that integrates sensory information and hormonal signaling, and is involved in the regulation of social behaviors. Lesion studies have shown a role for the MeA in copulation, most prominently in the promotion of ejaculation. The role of the MeA in sexual motivation, but also in temporal patterning of copulation, has not been extensively studied in rats. Here, we investigated the effect of chemogenetic inhibition and stimulation of the MeA on sexual incentive motivation and copulation in sexually experienced male rats. AAV5-CaMKIIa viral vectors coding for Gi, Gq, or no DREADDs (sham) were bilaterally infused into the MeA. Rats were assessed in the sexual incentive motivation test and copulation test upon systemic CNO or vehicle administration. We report that MeA stimulation and inhibition did not affect sexual incentive motivation. Moreover, both stimulation and inhibition of the MeA decreased the number of ejaculations in a 30 minute copulation test and increased ejaculation latency and the number of mounts and intromissions preceding ejaculation, while leaving the temporal pattern of copulation intact. These results indicate that the MeA may be involved in the processing of sensory feedback required to reach ejaculation threshold. The convergence of the behavioral effects of stimulating as well as inhibiting the MeA may reflect opposing behavioral control of specific neuronal populations within the MeA.

## 1. Introduction

Sexual behavior is an innately motivated behavior in the male rat and consists of three phases. During the initial phase, sexual incentive motivation propels a sexually experienced male into approach and investigation of a receptive female. After identification of the receptive female as a potential mate, the second phase of copulation quickly commences. Copulation consists of stereotypical motor output in the form of mounts and intromissions spaced over time in mount bouts, with chasing, genital grooming, and other non-copulation oriented behaviors in between. Multiple mounts and intromissions eventually culminate into ejaculation, the executive phase of sexual behavior. Even though there is no copulation without approach and no ejaculation without copulation, the behavioral output in different phases of sexual behavior might well be independently regulated on the neurobiological level (Ågmo 2002). This is supported by the notion that copulation parameters load onto different factors than anticipatory and approach parameters in factor analysis of male sexual behavior (Pfaus, Mendelson, and Phillips 1990). Studying the different phases of sexual behavior separately will lead to a more precise understanding of temporal and causal relations between neuronal activity and behavior.

The medial amygdala (MeA) is a sexually dimorphic brain region known to be involved in the regulation of a wide array of social behaviors, such as aggression, parental behavior, and sexual behavior, as reviewed in (Newman 1999; Sokolowski and Corbin 2012). These behaviors require the processing of contextual and sensory information in convergence with the internal state of the animal in order for the animal to display the appropriate behavioral response. Indeed, the high density of estrogen and androgen receptors, together with afferent input containing pheromonal and olfactory information, implicates the MeA as a primary locus for the integration of environmental and sensory information with the internal hormonal milieu of the animal (Swanson and Petrovich 1998; Simerly et al. 1990). Pheromonal information reaches the MeA directly from the accessory olfactory bulb, and olfactory information reaches the MeA from the main olfactory bulb via the cortical amygdala (Jennings and de Lecea 2020; Swanson and Petrovich 1998). Major efferent targets of the MeA include the medial preoptic area (mPOA), the bed nucleus of the stria terminalis, and the ventral medial hypothalamic nucleus (Canteras, Simerly, and Swanson 1995). These target areas have all been shown to be involved in the regulation of sexual behavior (Jennings and de Lecea 2020; Hull, Wood, and Mckenna 2006). The mPOA specifically is absolutely necessary for the display of sexual motivation and copulation in male rats (Hull, Wood, and Mckenna 2006). The MeA regulates dopamine release in the mPOA, and MeA lesion in addition to contralateral mPOA lesion is far more detrimental to copulation than MeA lesion in addition to ipsilateral lesion (Dominguez et al. 2001; Dominguez and Hull 2001; Kondo and Arai 1995). Considering its involvement in the processing of pheromonal and olfactory cues and its role as a major input area to the mPOA, implicates the MeA as a hub involved in the regulation of motivational, consummatory, and executive phases of sociosexual behaviors.

The role of the MeA in sexual motivation in rats has not been as extensively studied as its role in copulation. The involvement of the MeA in sexual approach would logically follow its integrative role for sensory information and hormonal signaling. Indeed, c-fos is induced in the MeA upon anogenital investigation or exposure to odors of receptive females in sexually experienced males (Baum and Everitt 1992). However, lesions of the MeA do not appear to affect incentive preference for an estrous female in male rats (Kondo and Sachs 2002), nor do they affect response latencies in a bar-pressing regimen in order to access an estrous female (Beck, Fonberg, and Korczyński 1982). In male hamsters, both the anterior and posterior MeA seem to be involved in the preference for odors of estrous females (Maras and Petrulis 2006). Further investigation of the role of the MeA in sexual incentive motivation is warranted.

The involvement of the MeA in the regulation of the copulatory phase of sexual behavior has been long established (Hull, Wood, and Mckenna 2006). This is apparent from data observing neuronal activity in the MeA during copulation and from studies manipulating the MeA during copulation. Single unit recordings in male rats show a remarkable increase in activity of MeA neurons upon the introduction of a receptive female (Minerbo et al. 1994). This activity remains high during the whole time period the receptive female is present and falls back down to baseline after removal of the female. In addition, neuronal activity spikes in the 20 seconds after copulation behaviors (a mount, intromission, or ejaculation). No increased neuronal activity was observed when a non-receptive female was introduced (Minerbo et al. 1994). In line with this, c-fos as well as Arc is induced upon copulation in the MeA of sexually experienced male rats (Turner et al. 2019; Coolen, Peters, and Veening 1997; Baum and Everitt 1992). Even though the MeA seems to clearly respond to copulation, different lesion studies in rats consistently find that the MeA is not essential for any aspect of copulation, including ejaculation (Harris and Sachs 1975; de Jonge et al. 1992; McGregor and Herbert 1992; Tsutsui, Shinoda, and Kondo 1994; Kondo, Sachs, and Sakuma 1997; Dominguez and Hull 2001). However, lesioning of the MeA does increase the ejaculation latency in behavioral tests (Harris and Sachs 1975; de Jonge et al. 1992; McGregor and Herbert 1992; Tsutsui, Shinoda, and Kondo 1994; Kondo, Sachs, and Sakuma 1997; Dominguez and Hull 2001). In addition, whereas the patterns of copulatory behavior look normal in MeA lesioned males, a larger number of mounts and intromissions usually precede ejaculation (Harris and Sachs 1975; Kondo and Arai 1995; Dominguez et al. 2001). Surprisingly, electrical stimulation of the MeA also dramatically impairs copulation (Stark et al. 1998). No further studies have investigated the effect of stimulating the MeA on sexual behavior in male rats. In all, these findings indicate a role for the MeA in copulation with regards to the processing of olfactory and pheromonal cues and somatosensory feedback from the penis, thereby affecting ejaculatory behavior.

Recently, progress has been made in the study of the role of the amygdala in sexual behavior of mice, where methodological advancements enabled a further interrogation of specific neuronal populations of the MeA. So far, studies that make use of more sophisticated techniques in rats are lacking. In addition, analysis of sexual behavior is often reduced to the annotation of only mounts, intromissions, and ejaculations. This prompted us to study the role of the MeA in sexual behavior in male rats by means of chemogenetics, with an extensive behavioral annotation allowing for additional analysis of temporal patterning of copulation through mount bout based assessment (Sachs and Barfield 1970). Because so little data exists on stimulation of the MeA in sexual behavior, we looked at the effects of both chemogenetic inhibition and stimulation of the MeA on sexual behavior in male rats. Importantly, with this study we assessed the involvement of the MeA in all stages of sexual behavior; sexual incentive motivation, and copulation (including ejaculation). We found that both stimulation and inhibition of the MeA disrupted ejaculation while increasing the number of copulatory behaviors preceding ejaculation, but did not affect sexual incentive motivation.

## 2. Materials and Methods

### 2.1 Animals

All rats (Charles River, Sulzfeld, Germany) were housed in Macrolon IV^®^ cages on a reversed 12h light/dark cycle (lights on between 23:00 and 11:00) in a room with controlled temperature (21 ± 1 °C) and humidity (55 ± 10%), with *ad libitum* access to standard rodent food and tap water. Rats were housed in same-sex pairs, unless otherwise noted (see brain surgery). In this experiment, 54 male Wistar rats were used as subjects. An additional 6 male Wistar rats were used as social incentives in the sexual incentive motivation (SIM) test. A total of 36 female Wistar rats were used as sexual incentives in the SIM test and as stimulus animals in the copulation test.

### 2.2 Viral constructs and drugs

Three viral constructs (University of North Carolina Vector Core, Chapel Hill, USA) were used in this experiment: AAV5-CaMKIIa-hM4D-mCherry (Gi), AAV5-CaMKIIa-hM3D-mCherry (Gq) and AAV5-CaMKIIa-EYFP (Sham). Clozapine N-oxide (CNO) (BML-NS105; Enzo Life Sciences, Farmingdale, USA) was dissolved in ddH_2_O at a stock concentration of 1 mg/mL (3 mM) and frozen at −20 °C in aliquots until further use. For experiments, rats were injected intraperitoneally with 1 mL/kg of the 1 mg/mL CNO solution or vehicle (ddH_2_O).

Silastic capsules (medical grade Silastic tubing, 0.0625 in. inner diameter, 0.125 in. outer diameter, Degania Silicone, Degania Bet, Israel) for females were 5 mm long and contained 10% 17β-estradiol (Sigma, St. Louis, USA) in cholesterol (Sigma, St. Louis, USA). The silastic tubing was closed off by inserting pieces of toothpick into both ends and sealed off with medical grade adhesive silicone (NuSil Silicone Technology, Carpinteria, USA).

Progesterone (Sigma, St. Louis, USA) was dissolved in peanut oil (Apotekproduksjon, Oslo, Norway) at a concentration of 5 mg/mL. Female rats were subcutaneously injected with 0.2 mL of the solution.

### 2.3 Surgical procedures

#### Ovariectomy

Stimulus females were ovariectomized under isoflurane anesthesia as previously described (Ågmo 1997). Briefly, a medial dorsal incision of the skin of about 1 cm was made, and the ovaries were located through a small incision in the muscle layer on each side. The ovaries were extirpated and a silastic capsule containing β-estradiol was placed subcutaneously through the same incision. The muscle layer was sutured and the skin was closed with a wound clip.

#### Brain surgery

Brain surgery consisted of subsequent bilateral infusions of the viral vector into the MeA. Rats were anesthetized with a mixture of zolazepam/tiletamine/xylazine/fentanyl (73.7 mg/73.7 mg/1.8 mg/10.3 μg per mL; 2 ml/kg) and placed in a stereotaxic apparatus (Stoelting Europe, Ireland). The skull was exposed through incision and small holes were drilled at the appropriate injection sites. A 30G cannula (Plastics One, Raonoke, USA) was inserted into each brain hemisphere sequentially at the following coordinates: AP −3,1 mm and ML ± 3,7 mm from bregma and DV −8,2 mm from the cortical surface (Paxinos and Watson 2007). Per infusion site, 750 nl of viral construct solution (Titers; Gi 4.3 × 10^12^ vg/mL, Gq 1.4 × 10^12^ vg/mL, Sham 7.4 × 10^12^ vg/mL) was injected at an infusion rate of 150 nl/min by a Hamilton syringe mounted in a minipump, connected to the infusion cannula by a piece of tubing (Plastics One, Roanoke, USA). Following infusion, the cannula was left in place for 10 minutes before withdrawal and closing of the skin with a continuous intradermal suture (Vicryl Rapide 4-0, Ethicon, Cincinnati, USA). After surgery, rats were single-housed for 3-7 days before being rehoused in pairs again. Analgesic treatment consisted of buprenorphine 0.05 mg/kg within 8 hours of surgery and every 12 hours for 72 hours thereafter.

### 2.4 Behavioral assessment

#### Sexual incentive motivation

The sexual incentive motivation test is described elsewhere (Ågmo 2003). Briefly, the SIM apparatus consists of a rectangular arena (100 × 50 × 45 cm) with rounded corners placed in a dimly lit (5 lx) room. At each long side, in opposite corners, a closed incentive stimulus cage was attached to the arena and separated from the arena by wire mesh (25 × 25 cm). A social stimulus (intact male rat) was placed in one of the stimulus cages and a sexual stimulus (receptive female rat) was placed in the other stimulus cage. To male subject rats, an intact male and a non-receptive female have the same salience as a social stimulus (Ågmo 2003). The subject rat was placed in the middle of the arena and video-tracked by Ethovision software (Noldus, Wageningen, the Netherlands) for 10 minutes. In Ethovision, virtual incentive zones (30 × 21 cm) were defined within the arena in front of each stimulus cage. The subject was considered to be within the zone whenever its point of gravity was. The software output consisted of the time the experimental subject spent in each incentive zone, the total distance moved, the time spent moving, and the mean velocity. From this data, the preference score was calculated (time spent in female incentive zones/total time in incentive zones). Subject rats were introduced right after each other, without cleaning the arena in between. The position of the stimulus cages (including the stimulus animal) was randomly changed throughout each experimental session. The SIM arena was cleaned with diluted acetic acid between experimental days.

#### Copulation

The copulation test was conducted in rectangular boxes (40 × 60 × 40 cm) with a Plexiglas front, in a room with lights on. During behavioral testing, the experimental subject was transferred from the room with the SIM test to the room with the copulation boxes. A receptive female was placed in the copulation box, after which the experimental subject was introduced. The test started upon introduction of the experimental subject male and lasted for 30 minutes. All test sessions were recorded on camera and behavior was later assessed from video. Behavioral assessment consisted of scoring behavioral events by means of the Observer XT software (Noldus, Wageningen, the Netherlands). For the entire 30 minutes, the copulatory behaviors mount, intromission and ejaculation were scored. During the first ejaculation series, we also behaviorally annotated 100% of the elapsed time by expanding the ethogram with clasping (mounting the female without pelvic thrusting), genital grooming (grooming of own genital region), other grooming (autogrooming in other regions than genital), chasing (running after the female), anogenital sniffing (sniffing the anogenital region of the female), head towards female (head oriented in the direction of the female while not engaging in other behavior), head not towards female (any behavior that is not oriented towards the female except grooming, such as walking, sniffing the floor, standing still with head direction away from female). From these data points the outcome measures as listed in table 1 were determined (see also (Heijkoop, Huijgens, and Snoeren 2018)). All behavioral tests were conducted during lights-off time.

**Table 1:** Copulation test outcome measure definitions

| Outcome measure | Definition |
| --- | --- |
| Number of ejaculations | Total number of ejaculations in the 30 min test |
| Latency to first ejaculation | Time from first copulatory behavior (mount or intromission) to ejaculation (NB: set to 1800 seconds in case no ejaculation was achieved during the test) |
| Latency to second ejaculation | Time from the end of the first post-ejaculatory interval to the next ejaculation |
| Mounts per ejaculation | Number of mounts in the first ejaculation series |
| Intromissions per ejaculation | Number of intromissions in the first ejaculation series |
| Intromission ratio | Number of intromissions in the first ejaculation series divided by the total number of copulatory behaviors in the first ejaculation series |
| Latency to first copulatory behavior | Time from the start of the test to the first copulatory behavior (mount or intromission) |
| Latency to first intromission | Time from the start of the test to the first intromission |
| Number of mount bouts per ejaculation | Number of mount bouts (a sequence of copulatory behaviors (one or more), uninterrupted by any behavior (other than genital autogrooming) that is not oriented towards the female) in the first ejaculation series |
| Mounts per mount bout | Mean number of mounts per mount bout in the first ejaculation series |
| Intromissions per mount bout | Mean number of intromissions per mount bout in the first ejaculation series |
| Inter-intromission interval | Time between intromissions in the first ejaculation series, calculated from the first intromission |
| Mount bout duration | Mean duration of mount bouts in the first ejaculation series |
| Time out duration | Mean duration of time-out (time from the end of one mount bout to the start of the next mount bout) |
| Post-ejaculatory interval | Time from the first ejaculation to the next copulatory behavior |
| Percentage of time spent on [behavior] | Percentage of time spent engaging in each of the annotated behaviors before the first ejaculation |
| Percentage of time spent in non-copulation oriented behavior | Percentage of time spent engaging in head not towards female + other grooming |

### 2.5 Brain processing, immunostaining and imaging

At the end of the experiment, rats were i.p. injected with a lethal dose of pentobarbital (100 mg/kg; Pentobarbital solution 100 mg/mL, Ås Produksjonslab AS, Ås, Norway) and, when deeply anesthetized, transcardially perfused with 0.1 M phosphate buffered saline (PBS; pH 7.4) followed by 4% formaldehyde in 0.1 M PBS. Brains were quickly removed and post-fixed in 4% formaldehyde in 0.1 M PBS for 48 hours. Subsequently, brains were transferred to a 20% sucrose in 0.1 M PBS solution, followed by a 30% sucrose in 0.1 M PBS solution until they had sunken. Brains were then either snap frozen by use of isopentane and kept at −80 °C until sectioning, or sectioned right away. Brains were sectioned on a cryostat (Leica CM1950, Leica Biosystems, Wetzlar, Germany, and Cryostar NX70, Thermo Fisher Scientific, Waltham, USA) into 30 μm thick sections and stored in cryoprotectant solution (30% sucrose w/v, 30% ethylene glycol v/v in 0.1 M phosphate buffer, pH 7.4) until further use.

For immunohistochemistry, 1 in every 5^th^ brain section within the area of interest was stained for the corresponding DREADD-conjugated fluorophore. For immunostaining, free-floating sections were washed in 0.1M Tris-buffered-saline (TBS, pH 7.6), blocked for 30 min in 0.5% BSA, and incubated on an orbital shaker for 24h at room temperature + 24h at 4 °C in polyclonal rabbit anti-mCherry (1:30 000, Abcam, cat. ab67453) or polyclonal chicken anti-EYFP (1:200 000, Abcam, cat. ab13970) antibody solution containing 0.1% Triton-X and 0.1% BSA in TBS. Sections were then incubated in biotinylated goat anti-rabbit (1:400, Abcam, cat. ab6720) or biotinylated goat anti-chicken (1:400, Abcam, cat. ab6876) antibody solution containing 0.1% BSA in TBS for 30 min, avidin-biotin-peroxidase complex (VECTASTAIN ABC-HRP kit, Vector laboratories, cat. PK-6100, dilution: 1 drop A + 1 drop B in 10 mL TBS) solution for 30 min, and 3,3’-diaminobenzidine solution (DAB substrate kit (HRP), Vector laboratories, cat. SK-4100, dilution: 1 drop R1 + 2 drops R2 + 1 drop R3 in 5 mL water) for 5 min, with TBS washes between all steps. Slides were dehydrated, cleared, and coverslipped using Entellan mounting medium (Sigma, St. Louis, USA).

After drying, the slides were loaded into an Olympus VS120 virtual slide microscope system. High resolution image scans were obtained for each section using a 20x objective (NA 0.75) and automatic focus and exposure settings in single plane. Using OlyVIA online database software (Olympus, Tokyo, Japan), viral spread was determined through assessment of the location and extent of stained cell bodies for separate brain regions. Animals that did not at a minimum have sufficient unilateral DREADD expression in the MeA were excluded from further analysis.

### 2.6 Design

Female stimulus animals were ovariectomized and implanted with a silastic capsule with β-estradiol at least one week before use in the SIM and copulation test. The females were injected with 1 mg progesterone 4 hours before use in behavioral tests in order to induce sexual receptivity.

Male subjects were first habituated to the SIM arena and sexually trained in three sessions over the course of a week. During the copulatory training sessions, that directly followed the SIM habituation, males were allowed to copulate with a receptive female in order to become sexually experienced. Males where then divided into three homogenous experimental groups based on the number of ejaculations in the last 30 minute copulation training session. Over the course of the second week, all male rats had brain surgery during which a viral vector carrying Gi(DREADD)-mCherry, Gq(DREADD)-mCherry, or EYFP genetic information, was infused bilaterally into the MeA. A 19-24 day recovery and DREADD expression period was allowed after surgery. Subsequently, rats underwent behavioral testing following an intraperitoneal injection of CNO and vehicle in a latin square within-subject design. Allowing a one week recovery period between copulation testing (enough for copulation parameters to return to baseline even after sexual exhaustion (Jackson and Dewsbury 1979)), each male was tested twice, once for each treatment, over the course of two weeks. Rats were first tested in the SIM test 30 min after i.p. injection with either vehicle or CNO. Following the SIM test, rats were tested in the copulation test 5-15 min later. Finally, rats were perfused with formaldehyde and brains were harvested for immunohistochemical analysis of DREADD expression.

The presented data in this manuscript consists of combined data from two separate homologous experiments.

### 2.7 Data analysis and statistics

Multiple linear mixed models employing virus as between-subject factor and treatment as within-subject factor were tested on the data using SPSS statistical software (IBM, version 26, Armonk, USA). Based on Akaike’s Information Criterion, a linear mixed model that included only the factors virus*treatment interaction term and experiment number as a covariate was deemed the best fit for the data. This mixed model was run for each of the separate outcome measures of the SIM test and the copulation test. In case of a significant virus*treatment interaction effect at the alpha 0.05 level, Bonferroni posthoc tests were conducted to identify significant within- and between-group differences. Supplementary analyses on small sample subgroups were done by employing t-tests (alpha 0.05) without multiple comparison correction.

The SIM preference score was compared to chance (0.5) with a one-sample t-test for each treatment within each group. Time spent in female zone was compared to time spent in male zone for each treatment within each group with a paired t-test. For an effect on sexual incentive motivation, comparisons between both the preference scores and the time spent in female zone needs to be statistically significant, as an increased preference score is irrelevant when the total time spent in incentive zones is relatively small.

## 3. Results

### 3.1 DREADD expression

DREADD expression in the MeA was assessed by immunohistochemical staining. Out of 56 animals, 2 animals died before perfusion and were excluded because of a lack of histological data. Another 11 animals were excluded due to insufficient MeA DREADD expression.

Somatic DREADD expression in the remaining 43 animals was observed throughout the anterior and posterior MeA, with higher density posteriorly (Figure 1). In the majority of animals, DREADD expression extended to amygdaloid structures lateral and posterior from the MeA, namely the intraamygdaloid division of the bed nucleus of the stria terminalis (STIA), the amygdalohippocampal area (Ahi), the posteromedial cortical nucleus (PMCo), and the basomedial amygdaloid nucleus (BM). Most animals also had low-density ventral hippocampal (vHC) DREADD expression. In addition, low-density, mostly unilateral, expression was observed in the peduncular part of the lateral hypothalamus (LH) in 16 animals.

**Figure 1.**
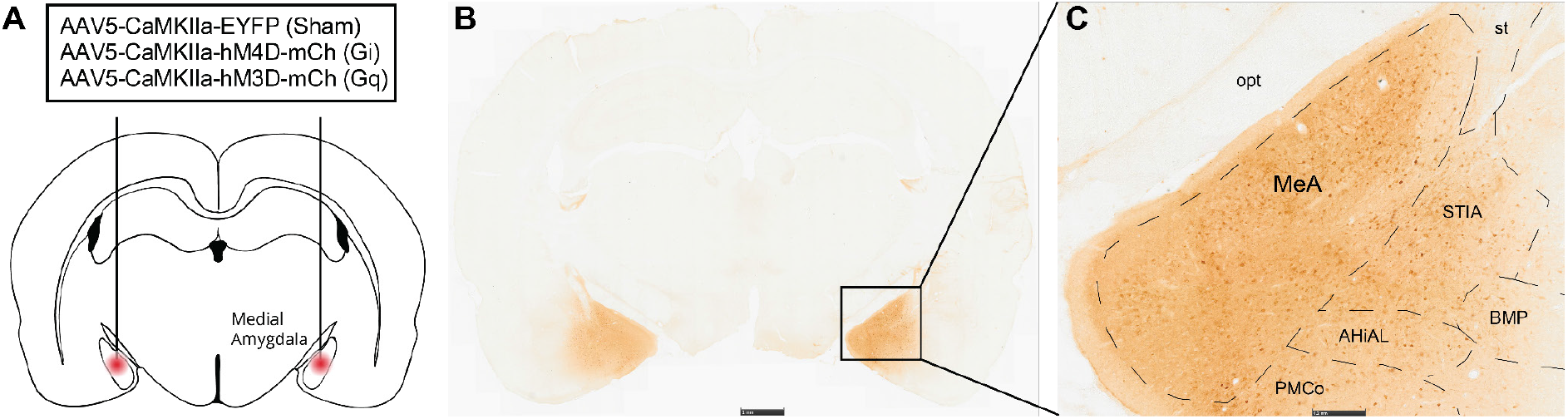
Medial amygdala DREADD expression. **(A)** Bilateral viral targeting of the MeA. **(B)** Example DREADD expression on whole brain section at approximately AP −3.2 from bregma. **(C)** Magnified inset of (B) showing somatic DREADD expression in the MeA and surrounding structures. MeA = medial amygdala; opt = optic tract; st = stria terminalis; STIA = intraamygdaloid division of the bed nucleus of the stria terminalis; BMP = basomedial amygdaloid nucleus, posterior part; AHiAL = amygdalohippocampal area, anterolateral part; PMCo = posteromedial cortical amygdaloid nucleus.

### 3.2 Sexual incentive motivation

To study the involvement of the MeA in sexual incentive motivation, we compared SIM test (Fig. 2A) parameters in vehicle (VEH) and CNO treated Sham, Gi-DREADD, and Gq-DREADD males. Subject males in each virus group (Sham, Gi, and Gq), and during each treatment, significantly spent more time in the female zone compared to the male zone (Fig. 2C; Sham-CNO t(15)=13.7, Sham-VEH t(15)=13.6, Gi-CNO t(11)=15.8, Gi-VEH t(11)=10.8, Gq-CNO t(14)=11.3, Gq-VEH t(14)=9.45, p<0.001 for all groups). This was also reflected in the preference scores that were significantly larger than 0.5 (Fig. 2D; Sham-CNO t(15)=14.6, Sham-VEH t(15)=18.9, Gi-CNO t(11)=19.4, Gi-VEH t(11)=12.6, Gq-CNO t(14)=12.6, Gq-VEH t(14)=11.2, p<0.001 for all groups). Additionally, subject males visited the female zone more frequently in all but Gi-CNO (Suppl. Fig 1A; Sham-CNO t(15)=3.41, p=0.004, Sham-VEH t(15)=4.25, p<0.001, Gi-VEH t(11)=3.53, p=0.005, Gq-CNO t(14)=2.63, p=0.020, Gq-VEH t(14)=2.86, p=0.013). There was a shorter latency to visit the female zone than to visit the male zone in Gi-CNO (Suppl. Fig 1B; t(11)=2.27, p=0.044) and Gi-VEH (Suppl. Fig 1B; t(11)=2.45, p=0.032). We found no significant interactions of treatment and virus for distance moved (Figure 2B), time spent in zones (Fig. 2C), preference score (Fig. 2D), frequency of zone entry (Suppl. Fig 1A), latency to enter zone (Suppl. Fig 1B), time spent moving (Suppl. Fig 1C), and mean velocity (Suppl. Fig. 1D).

**Figure 2.**
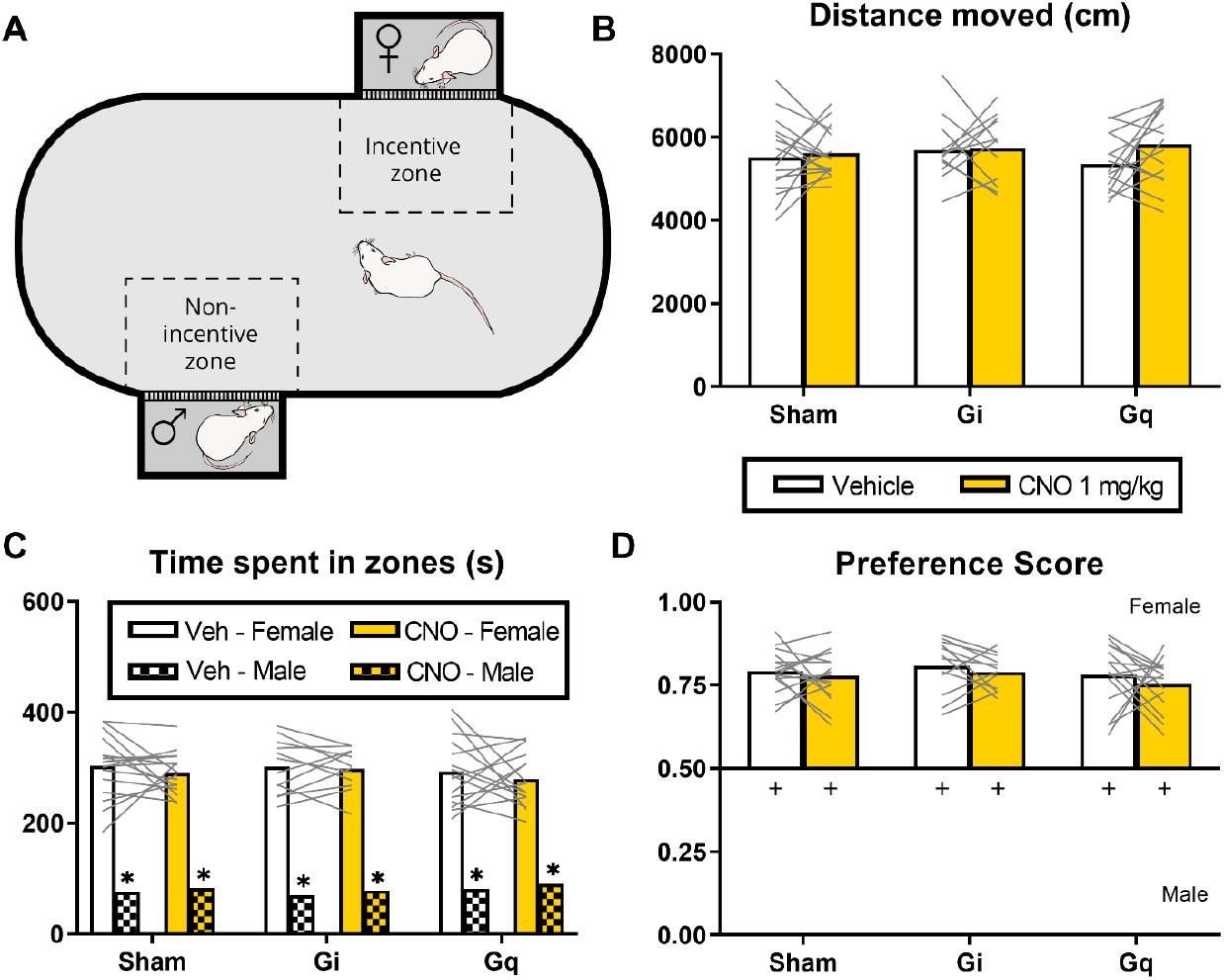
Silencing or stimulating the MeA does not affect sexual incentive motivation. **(A)** Sexual incentive motivation test (10 min). **(B)** Total distance moved during the 10 minute test. **(C)** Total time spent in the incentive zone (female zone) and the non-incentive zone (male zone). *p<0.05 compared to “female zone” **(D)** Preference score (time spent in female zone/total time spent in female and male zones). ^+^p<0.05 compared to 0.5. **All panels:** n = 16 (sham), 12 (Gi), 15 (Gq).

### 3.3 Copulation

Immediately after the SIM test, male subjects were tested in the copulation test (Fig. 3A). No effects of MeA silencing or stimulation on latency to first copulatory behavior (Fig. 3B), nor on latency to first intromission were observed (Suppl. Fig. 2). We did find that ejaculation parameters were significantly affected (Fig. 3C). CNO decreased the number of ejaculations during the 30 minute test (Fig. 3C; virus x treatment: F(5,44)=11.28, p<0.001) in both the Gi-group (Mean difference (md)=1.08, p<0.001, *g*=0.83) and the Gq-group (md=1.27, p<0.001, *g*=1.267) compared to vehicle. Although, only in Gi-CNO were the number of ejaculations also significantly decreased (md=1.22, p=0.011, *g*=1.23) compared to Sham-CNO. The decrease in number of ejaculations logically followed a significant CNO-induced increase of latency to ejaculation (Fig. 3C; virus x treatment: F(5,47)=6.58, p<0.001) in both the Gi-group (md=490, p<0.001, *g*=0.91) and the Gq-group (md=380, p=0.001, *g*=0.83) compared to vehicle, and only in Gi-CNO compared to Sham-CNO (md=481, p=0.009, *g*=1.13). These effects persisted during the second ejaculation series (Fig. 3C; virus x treatment: F(5,32)=4.890, p=0.002). CNO increased the latency to second ejaculation compared to vehicle in the Gi-group (md=162, p=0.003, *g*=0.53), as well as in the Gq-group (md=162, p=0.003, *g*=1.35), and compared to Sham-CNO in the Gi-group only (md=195, p=0.009, *g*=0.98). Further analysis of the first ejaculation series showed that CNO significantly increased the number of mounts compared to vehicle (Fig. 3D; virus x treatment: F(5,51)=2.41, p=0.049) in both the Gi-group (md=13.5, p=0.012, *g*=0.86) and the Gq-group (md=10.4, p=0.029, *g*=0.76). The number of intromissions preceding the first ejaculation was also affected by CNO compared to vehicle in the Gi-group (Fig. 3D; virus x treatment: F(5,48)=4.63, p=0.002; Gi md=8.5, p=0.001, *g*=0.83), as well as in the Gq-group (md=6.27, p=0.005, *g*=0.96). However, no statistical significant effects were observed on the number of mounts and intromissions between Gi-CNO or Gq-CNO compared to Sham-CNO. The numbers of mounts and intromissions were proportionally increased by CNO in the Gi- and Gq-groups compared to vehicle, as intromission ratio remained unaffected by CNO in both these groups (Fig. 3E). The larger number of copulatory behaviors did not lead to an increase in the mean number of mounts and intromissions per mount bout, nor the mean duration of mount bouts (Suppl. Fig. 2). Instead, it was reflected in a CNO-induced increase of the number of mount bouts preceding ejaculation (Fig. 3F; virus x treatment: F(5,49)=5.55, p<0.001) in both the Gi-group (md=18.6, p<0.001, *g*=0.95) and the Gq-group (md=13.9, p=0.002, *g*=01.25) compared to vehicle. But again, there was no statistical significant effect between Gi-CNO or Gq-CNO compared to Sham-CNO. Finally, no effects were observed on parameters of temporal patterning; mean duration of time-out (Fig. 3G), post-ejaculatory interval (Fig. 3H), and inter-intromission interval (Suppl. Fig. 2).

**Figure 3.**
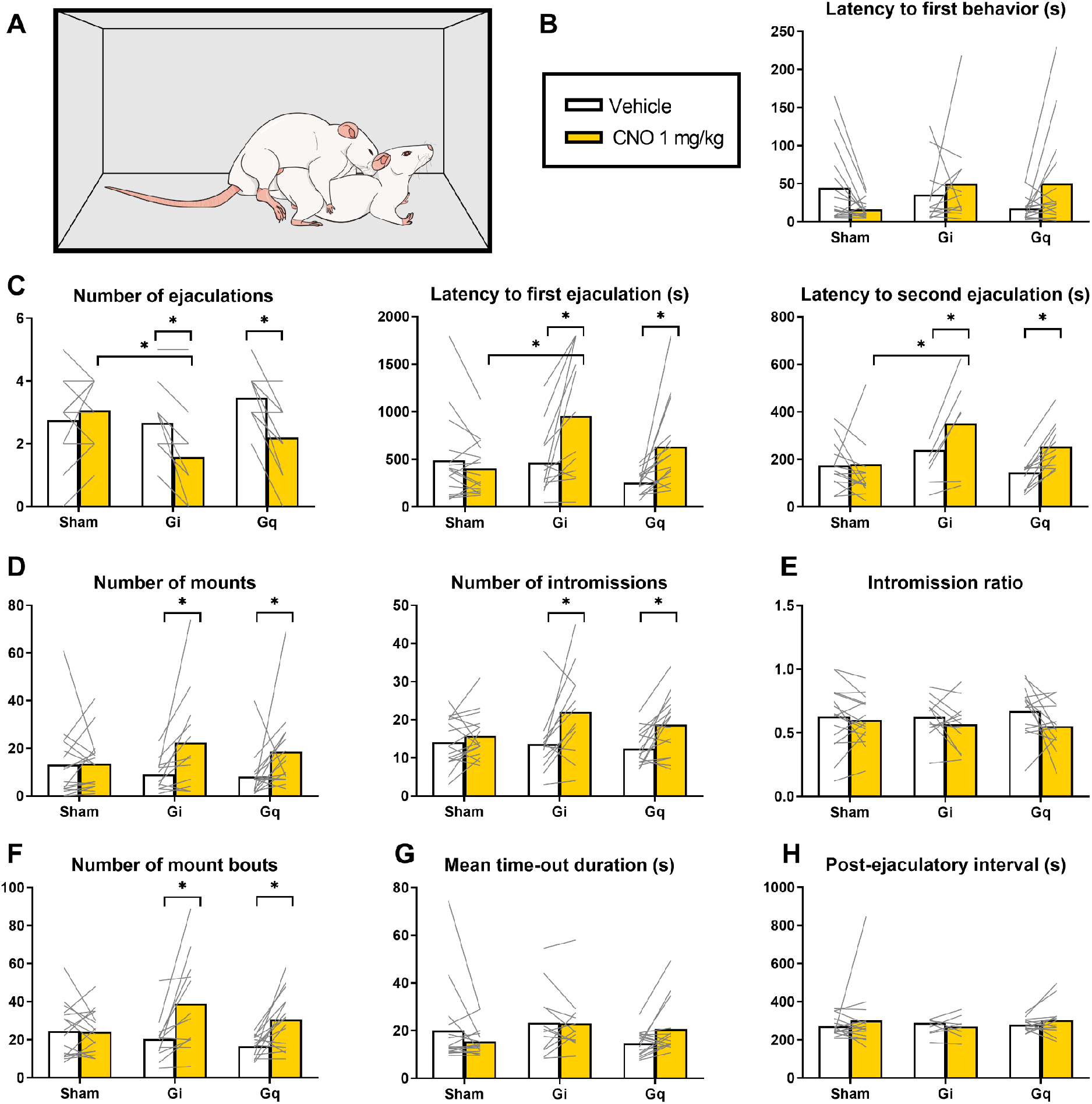
Silencing and stimulating the MeA affect copulation parameters in the same direction. **(A)** Copulation test (30 min). **(B)** Latency to first copulatory behavior, i.e. mount or intromission. **(C)** Ejaculation parameters: Number of ejaculations, Latency to first ejaculation, and Latency to second ejaculation (n = 13 (sham), 6 (Gi), 12 (Gq)). **(D)** Number of mounts and number of intromissions in the first ejaculation series. **(E)** Intromission ratio (intromissions/(mounts+intromissions)) in the first ejaculation series. **(F)** Number of mount bouts (one or more uninterrupted copulatory behaviors) in the first ejaculation series. **(G)** Mean duration of time-outs (intervals between mount bouts) in the first ejaculation series. **(H)** Post-ejaculatory interval of the first ejaculation series (n = 15 (sham), 8 (Gi), 14 (Gq)). **All panels:** *p<0.05; n = 16 (sham), 12 (Gi), 15 (Gq) unless otherwise indicated.

Analysis of the percentage of time spent on each of the behavioral parameters showed significant effects of CNO on the percentage of time spent on head not towards female (Suppl. Fig. 3; virus x treatment: F(5,45)=3.37, p=0.011) compared to vehicle within the Gq-group only (md=8.01, p=0.009), but not for Gq-CNO vs. Sham-CNO. Consequently, a statistical significant effect was found for percentage of time spent on non-copulation oriented behavior (Suppl. Fig. 3; virus x treatment: F(5,44)=4.08, p=0.004), which is comprised of percentage of time spent on head not towards female and other grooming, for Gq-CNO compared to Gq-vehicle (md=9.14, p=0.004), but not for Gq-CNO compared to Gq-vehicle. No significant interaction of virus and treatment was found in percentage of time spent on other grooming, genital grooming, anogenital sniffing, chasing and clasping (Suppl. Fig. 3).

## 4. Discussion

The MeA is a sexually dimorphic brain region involved in the regulation of sexual behavior (Hull, Wood, and Mckenna 2006; Newman 1999). The afferent and efferent connections of the MeA and the expression of hormonal receptors and aromatase in the MeA suggest its involvement in integrating environmental and sensory information with the internal hormonal state of the animal (Swanson and Petrovich 1998; Canteras, Simerly, and Swanson 1995; Simerly et al. 1990; Jennings and de Lecea 2020). Considering the position of the MeA as an important integration area, and input area of the mPOA, we aimed to shine more light on the role of the MeA during all stages of sexual behavior in male rats. Our main finding here was that both silencing and stimulating the MeA did not impair incentive motivation or alter the structure and patterns of copulatory behavior, but did result in increased ejaculation latency and consequently a decrease in the number of achieved ejaculations during a 30 minute test.

Our findings were in line with MeA lesion studies (Harris and Sachs 1975; de Jonge et al. 1992; McGregor and Herbert 1992; Tsutsui, Shinoda, and Kondo 1994; Kondo, Sachs, and Sakuma 1997; Dominguez and Hull 2001), as we found that silencing of the MeA impaired ejaculation as shown by an increased latency to ejaculation, and consequently also caused a reduction in the achieved number of ejaculations. Similar to what others found (Harris and Sachs 1975; Kondo and Arai 1995; Dominguez et al. 2001), we also observed that more mounts and intromissions preceded ejaculation, while the intromission ratio was not affected. This indicates that erectile function is not impaired by MeA silencing. In our more extensive behavioral analysis, we annotated 100% of the time until the second ejaculation series. This allowed the assessment of temporal patterning of copulation by further analysis of mount bouts and time-outs (Sachs and Barfield 1970). We showed that the temporal pattern of copulation remained unaffected by silencing of the MeA. Together, these findings lead us to infer that the increased ejaculation latency is not caused by a decreased erectile function or a decreased copulatory pace, but may rather be attributable to a decreased sensitivity to penile stimulation. This is congruent with findings that show that in males, c-fos in the MeA is induced upon penile stimulation (intromissions and ejaculations) (Baum and Everitt 1992; Veening and Coolen 1998), and in females upon vaginal-cervical stimulation (Tetel, Getzinger, and Blaustein 1993), indicating a role for the MeA in the processing of sensory information. Interestingly, c-fos in the MeA upon ejaculation is expressed in a cell cluster more lateral in the MeA, whereas c-fos expression upon copulation and odor exposure is more diffusely located medially in the MeA (Coolen, Peters, and Veening 1997; Veening and Coolen 1998). The activity of the specific subset of lateral neurons associated with ejaculation could mean that these neurons respond to the sensory signal of ejaculation, or it could mean that they are involved in the actual orchestration of ejaculation. Our study shows that chemogenetic manipulation of the MeA impaired ejaculation, showing a role for the MeA in the relay of information that leads to the orchestration of ejaculation. Thus, the processing and accumulation of sensory feedback may occur in the MeA, which ultimately leads to the reach of ejaculation threshold.

Surprisingly, we found the same, attenuated, effects on copulation when stimulating the MeA as when inhibiting the MeA, although only the Gi-group reached statistical significance when comparing ejaculatory parameters to the Sham-group. These findings correspond to a study by Stark et al. (Stark et al. 1998), who found that electrical stimulation of the MeA in sexually experienced male rats reduced chasing, sniffing, and mounting of an estrous female while it increased these behaviors towards a non-estrous female (Stark et al. 1998). The authors hypothesized that the increased mounting of a non-estrous female may actually reflect an increase in aggressive behavior caused by MeA stimulation, which would be suppressed by the sensory cues emitted by an estrous female. Some recent studies in mice might provide an explanation for these findings. It was demonstrated that high laser intensity optogenetic stimulation of all neurons or GABAergic neurons selectively in the MeA leads to aggression towards both male and female intruders, whereas low laser intensity (with same frequency and pulse duration) optogenetic stimulation of GABAergic neurons triggers anogenital sniffing and mounting (Hong, Kim, and Anderson 2014). A similar scalable behavioral control by laser intensity was found in Esr1+ cells in the mouse ventromedial hypothalamus (Lee et al. 2014). It was demonstrated in this latter study that higher laser power both activates more neurons, as well as increases the average activity per neuron. In addition, chemogenetic activation of glutamatergic neurons in the MeA suppressed all social behavior and promoted self-grooming in mice (Hong, Kim, and Anderson 2014). Next to that, a large proportion of neurons in the MeA respond preferentially to one sex of conspecifics (Bergan, Ben-Shaul, and Dulac 2014), indicating a role for the MeA to identify an appropriate mate and assure the appropriate behavioral response. Thus, a model could be proposed in which different neuronal populations in the MeA, with different activation thresholds, might orchestrate either sexual behavior or aggression or attenuate social behaviors in general, depending on the sensory cues emitted by the conspecific stimulus animal. We observed no aggression or reduced chasing, sniffing, and mounting in any of our subject males towards estrous females upon MeA stimulation, but stimulatory properties of electrical probes, optogenetics and chemogenetics are different in nature. Where effects of electrical and optogenetical stimulation are dependent on the voltage/laser power, and stimulation frequency applied, it is not possible to modulate stimulatory properties of chemogenetic stimulation. If aggressive and copulatory behavioral output in rats is dependent on the intensity of MeA stimulation as it is in mice, the electrical stimulation by Stark et al. and the chemogenetic stimulation in our study, with extensive DREADD expression, could theoretically have been “out of range” for observations of stimulatory effects on ejaculation or copulatory pace. More importantly, we did not target any specific neuronal subpopulation in our study, and so opposing effects of manipulation of GABA-ergic (inter)neurons and glutamatergic neurons in the MeA could have led to a diffuse effect of chemogenetic stimulation as well as inhibition. Whether neuronal subpopulations in the MeA of male rats have similar opposing effects on sexual behavior as in mice remains to be investigated.

In the current study, we employed a more extensive analysis of temporal patterning of copulation. Sachs and Barfield showed that male rats copulate in mount bouts (uninterrupted sequence of mounts and/or intromissions) and that the intervals between these mount bouts (time-outs) are highly constant (Sachs and Barfield 1970). Mount bouts are not intromission driven, and copulatory pace is therefore better expressed in the time-out duration than in the inter-intromission interval. Our mount bout analysis here allowed us to conclude that even though males took longer to ejaculate, copulatory behavior patterns remained unaffected, as was reflected in unaffected mount bout structure (mounts and intromissions per mount bout) and interval durations (time outs). Mount bout analysis provides valuable insight in assessment of sexual behavior of male rats and we stress that it should be part of future studies employing behavioral annotation of copulation.

Silencing and stimulation of the MeA did not interfere with the preference for an estrous female over a social stimulus. In a study by Kondo and Sachs (Kondo and Sachs 2002), small lesions of the posterior MeA also did not affect preference for an estrous female over a non-estrous female in a similar set-up as ours, albeit with the females being behind opaque walls preventing visual cues to the subject male. In the same study it was found that the preference for an estrous female was attenuated in MeA-lesioned males compared to sham-lesioned control males if the stimulus females were anesthetized. In this set-up, the only sensory modalities available to the subject animal would have been audition and olfaction, which is not sufficient to induce preference over a social stimulus in male rats (Ågmo and Snoeren 2017). These results of these studies imply that olfaction-induced sexual approach is reliant on the MeA, but that the processing of this information is not necessary to maintain sexual incentive motivation and preference when multiple sensory modalities are present. Interestingly, unconditioned pre-exposure to an inaccessible estrous female decreases ejaculation latency in sexually experienced males, but not in naïve males, in a directly following copulation test, and this effect is blocked by lesions of the MeA (de Jonge et al. 1992). In addition, chemogenetic silencing of the MeA attenuated male urine odor preference in sexually naïve female mice (McCarthy et al. 2017). Together with the notion that MeA lesions almost completely block copulation in sexually naïve male rats (Kondo 1992), a far larger effect than in sexually experienced animals, this emphasizes how experience shapes the role of the MeA in different aspects of sexual behavior. Therefore, it could well be that chemogenetic stimulation and inhibition of the MeA of sexually naïve males would result in different findings, even though the fact that we did not find any effects on sexual incentive motivation is in line with the possibility that sexual approach and copulation may rely on different neurobiological mechanisms (Ågmo 2002). Finally, it should be noted that specific neuronal populations in the MeA have been shown to be involved in sexual approach behavior in mice, and that our null-findings could also be a result of non-specific targeting diffusing opposing effects (Yao et al. 2017; Adekunbi et al. 2018).

A limitation of our study is that some of the subject males in our study had DREADD expression in the lateral hypothalamus, a brain area known to be involved in sexual behavior, specifically ejaculation, the post-ejaculatory interval, and preference for an estrous female (Lorrain et al. 1997; Singh, Desiraju, and Raju 1996; Kippin et al. 2004). We ran a sub-analysis on our data set excluding all animals with LH expression, and this resulted in similar findings. The expression of DREADD also extended to structures outside of the MeA in this study. The majority of animals expressed DREADD in the STIA, AHi, PMCo, and BM at a similar density as in the MeA, and some animals had low density expression in the vHC as well. Some of the amygdaloid nuclei expressing DREADD have been implicated in the regulation of aspects of sexual behavior (Kondo 1992; Coolen, Peters, and Veening 1997; Romero, Beltramino, and Carrer 1990; Maras and Petrulis 2008; Root et al. 2014; Yamaguchi et al. 2020). In an additional analysis of a subset of a few animals that solely and substantially expressed DREADD in structures posterior from the MeA (i.e. AHi, PMCo, BM, and vHC), we found no indication of any effects on sexual incentive motivation or copulation. Even though we cannot be completely certain that the DREADD-expressing brain areas outside of the MeA did not contribute to the measured effects in our data set, we conclude that the main effects that we found are attributable to manipulation of the MeA.

Integrating our results on sexual incentive motivation and copulation with the literature suggests that the MeA has a role in the processing of sexually arousing stimuli in male rats before and during copulation. We hypothesize that even though cue processing by the MeA before the start of copulation may not influence the incentive preference for an estrous female in the presence of all sensory modalities, it might rather impact the state of arousal during subsequent copulation, an effect shaped by sexual experience. Our current experimental design did not allow for exploration of this hypothesis, which should be further assessed in future research. Our study showed that the MeA is involved in the regulation of ejaculation. The increased latency to ejaculation is not caused by effects on temporal patterning of copulation or erectile function. Rather, we conclude that the MeA has a role in the processing of sensory feedback necessary to overcome ejaculation threshold during copulation.

## Acknowledgements

Financial support was received from Norwegian Research Council; grant #251320 to EMS. We thank Carina Sørensen, Katrine Harjo, Ragnhild Osnes, Remi Osnes and Nina Løvhaug for the excellent care of the animals. All imaging was conducted by use of equipment at the Advanced Microscopy Core Facility at UiT The Arctic University of Norway.

## Author contributions

PTH: Experimental design, carrying out experiment, methodology, analysis, writing - original draft

RH: Experimental design, methodology, supervision, writing – review and editing

EMS: Experimental design, methodology, supervision, writing – review and editing, funding acquisition

## Supplementary figures

**Supplementary figure 1.**
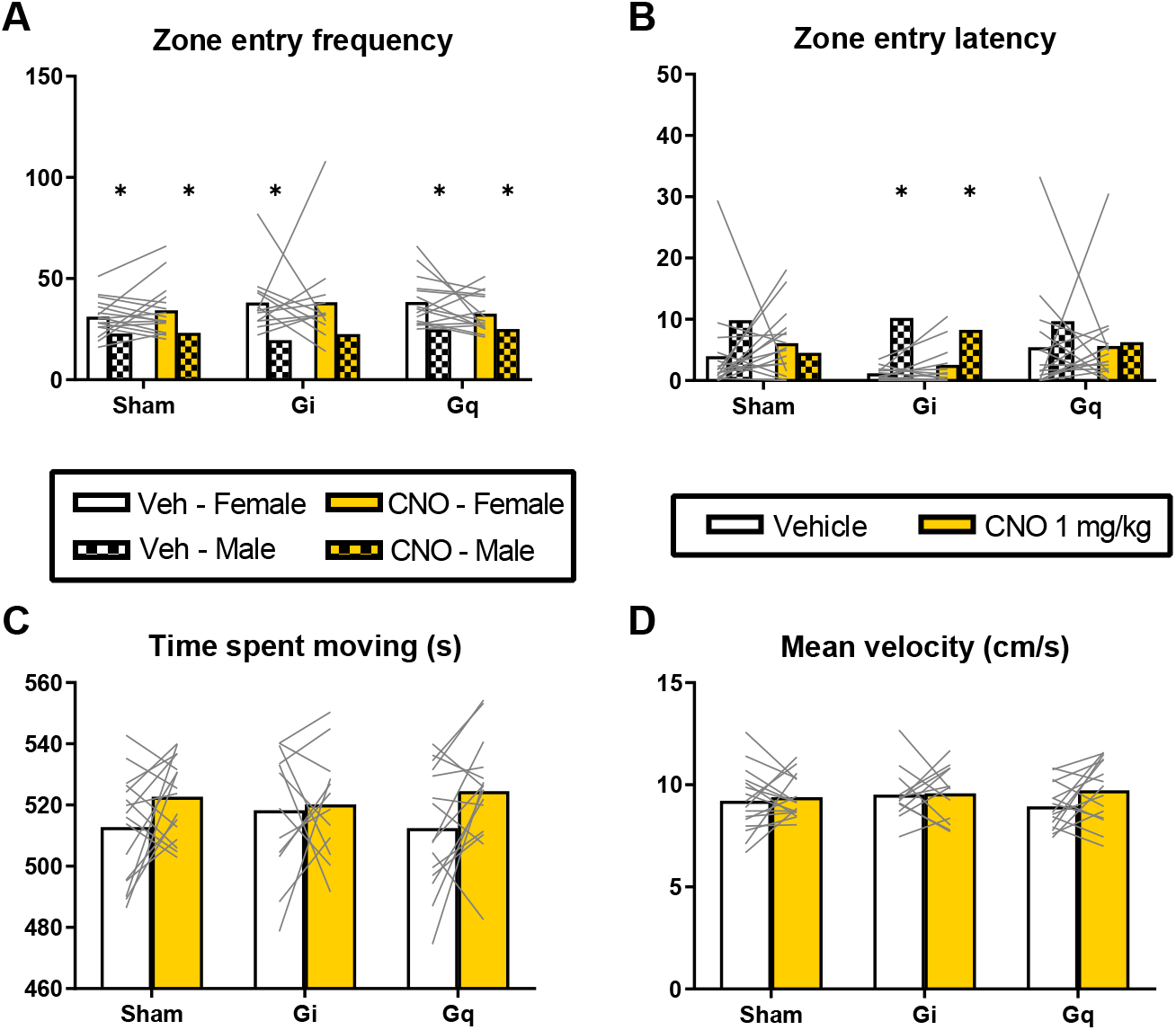
Additional SIM test outcome measures. Zone entry frequency. **(B)** Zone entry latency. **(C)** Time spent moving. **(D)** Mean velocity. **All panels:** *p<0.05 compared to “female zone”.

**Supplementary figure 2.**
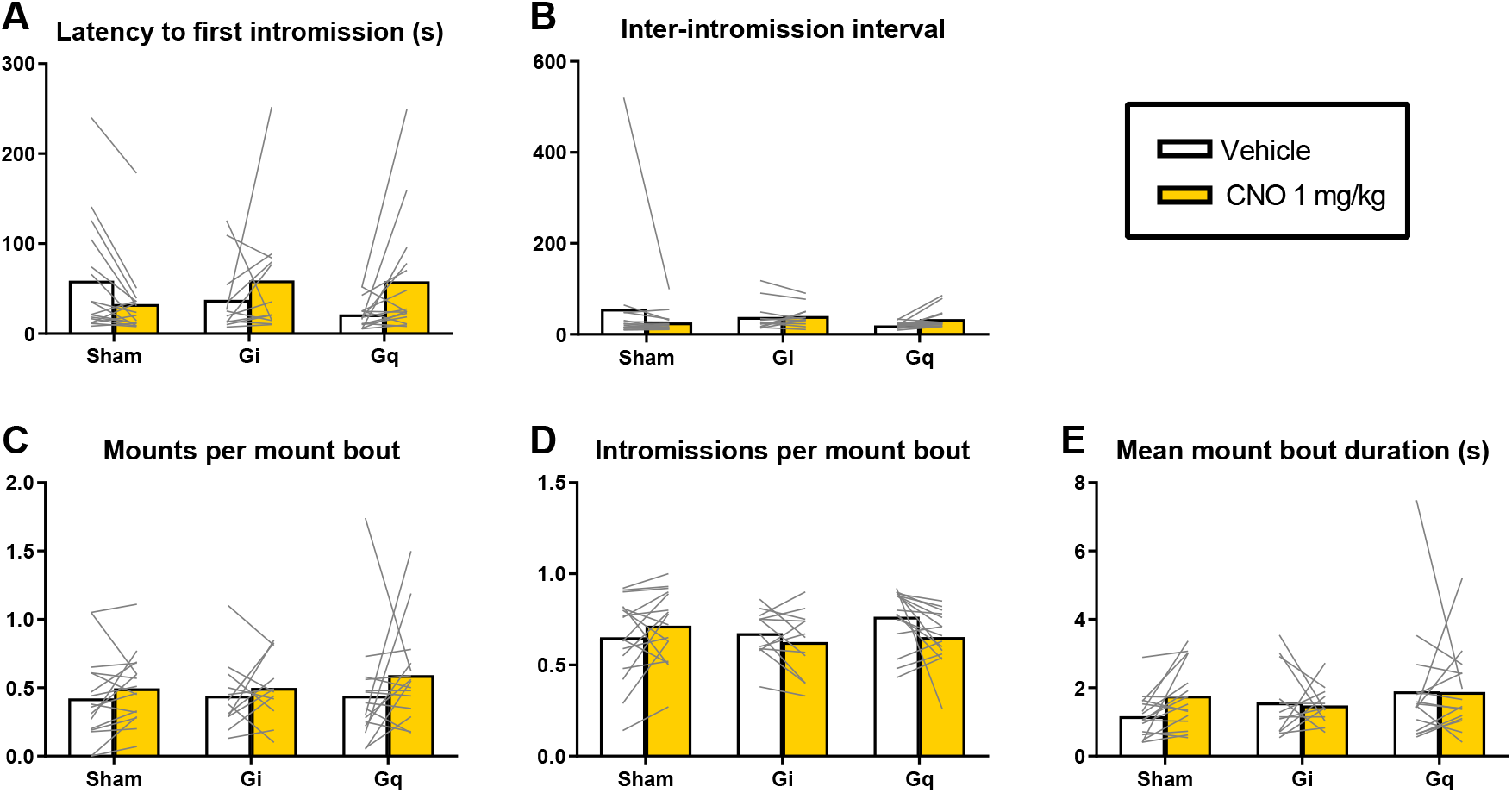
Additional copulation test outcome measures (first ejaculation series). **(A)** Latency to first intromission. **(B)** Inter-intromission interval. **(C)** Mounts per mount bout. **(D)** Intromissions per mount bout **(E)** Mean mount bout duration.

**Supplementary figure 3.**
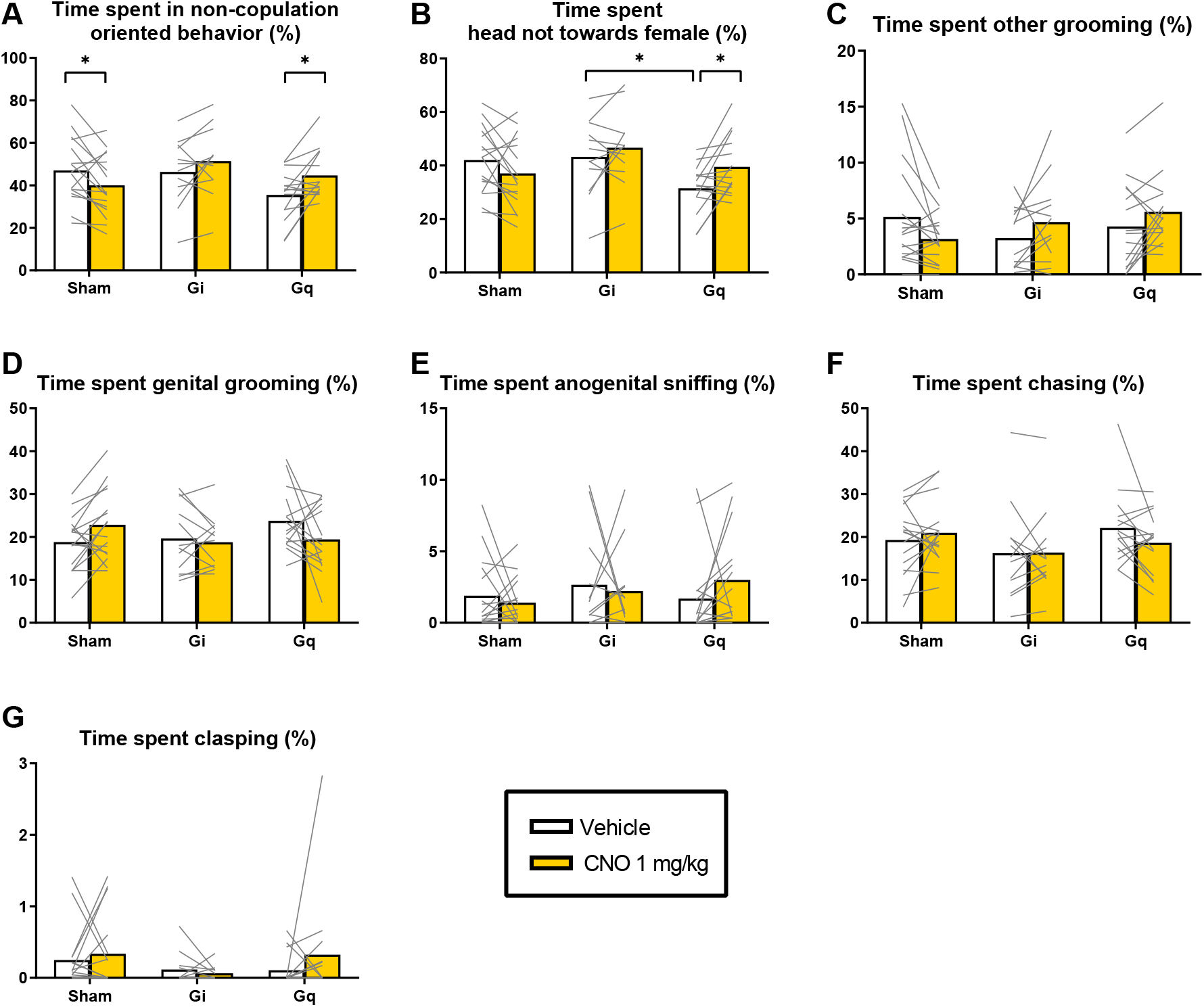
Percentage of time spent on behaviors during first ejaculation series. **(A)** Non-copulation oriented behavior (other grooming + head not towards female). **(B)** Head not towards female. **(C)** Other grooming. **(D)** Genital grooming. **(E)** Anogenital sniffing. **(F)** Chasing. **(G)** Clasping. **All panels:** *p<0.05

